# Whole-Genome Sequencing and Phylogenetic Analysis of *Anolis adenovirus 2* Reveals Conserved Genome Organization and Gene-Specific Evolutionary Patterns

**DOI:** 10.64898/2026.07.23.740413

**Authors:** Cleo H. Falvey, Anthony J. Geneva

## Abstract

Adenoviruses, which infect vertebrates, have a rich history of evolution that includes both host switching and coevolution, particularly within *Barthadenovirus*, a genus that infects squamate reptiles, birds, and mammals. Potential host-switching events can be identified by comparing the evolutionary histories between viruses and their hosts; however, many *Barthadenovirus* phylogenies have been inferred based on a limited number of easily-amplifiable gene segments. Whole-genome sequencing novel strains of *Barthadenovirus* can provide greater phylogenetic confidence, and therefore more accurately identify host-switching when it occurs. Here, we present the whole-genome sequence, annotation, reconciled species tree, and molecular evolution analyses of two isolates of *Anolis adenovirus 2*, a member of *Barthadenovirus*. Our two isolates of *Anolis adenovirus 2* are sister lineages with very high sequence similarity. Our results support existing hypotheses regarding the ancestral hosts of *Barthadenovirus* (squamate reptiles), and proposed host switching events within and between squamate reptiles and other vertebrate classes. We leverage our novel genome annotations to perform comparative synteny analyses, identifying a set of shared genes across *Barthadenovirus* whose gene order is largely conserved within the genus. Finally, our molecular evolution analyses highlight trends in evolutionary pressures on individual genes: Genes associated with viral replication and structure have experienced slower rates of evolution than those encoding proteins involved in host interaction. Our two sequenced isolates of *Anolis adenovirus 2* add to an expanding number of *Adenovirus* genomic resources and facilitate future investigations into the patterns and processes shaping adenovirus diversification.

## 1 Introduction

Sequencing the genomes of viruses belonging to families that are capable of host-switching, or jumping between different host species, can elucidate the evolutionary patterns underlying these processes and make predictions about the mechanisms that may enable host-switching (Longdon et al., 2014; Kaján et al., 2020; Borkenhagen et al., 2019). *Adenoviruses* are a diverse group of viruses that infects organisms across the vertebrate tree of life. There are six genera currently recognized: *Mastadenovirus*, *Aviadenovirus*, *Ichtadenovirus*, *Siadenovirus*, *Testadenovirus*, and *Barthadenovirus*. *Barthadenoviruses* (previously *Atadenovirus)* have been found to infect diverse species of squamate reptiles (Hyndman et al., 2019; Julian and Durham, 1982; Ball et al., 2014b; Farkas et al., 2008; Prado-Irwin et al., 2018; Ascher et al., 2013), ungulates (Lung et al., 2022; Jesse et al., 2022; Harrach, 2000), and birds (Matsvay et al., 2022; Bashashati et al., 2023). The evolutionary history of *Barthadenovirus* and the phylogenetic diversity of its hosts suggest that there have been ancestral host-switching events across this clade, likely from squamates into ruminants and birds, making it an excellent model for the study of viral evolution (Harrach, 2000; Harrach et al., 2019; Marschang, 2011). Studying viruses that infect hosts with largely complete and well-resolved phylogenies can enable more accurate reconstruction of host-virus coevolutionary relationships (Kaján et al., 2020). Fortunately, *Barthadenovirus* has been found in *Anolis* lizards, which are a well-established model organism for the study of evolution, and have a robust, well-established phylogeny with more than 400 species (Losos, 2009; Poe et al., 2017; Román-Palacios et al., 2018).

Previous research has identified *Barthadenovirus* infection in several species of *Anolis* lizards, including one strain in *Anolis carolinensis* (Ball et al., 2014a), eight diverse strains in *Anolis sagrei* (Prado-Irwin et al., 2018) that showed signatures of both coevolution and host-switching, and two subspecies of *Anolis distichus*: Anolis adenovirus 1 in *Anolis distichus ignigularis* and Anolis adenovirus 2 in *Anolis distichus ravitergum* (Ascher et al., 2013). These studies are limited, however, by the fact that these phylogenies were based on a conserved 300 base-pair long segment of the DNA polymerase gene (DPOL). This relatively short region was selected as the focal point of previous analyses due to its presence in all known *Adenovirus* strains and because primers designed to reliably amplify this region have been used widely, generating a large public dataset of sequences representing most known *Adenovirus* strains (Wellehan et al., 2004a). While many useful insights have come from analyzing this gene, sequencing the whole genome of the virus will allow us to gain deeper insight into the evolutionary history of these viruses by allowing for the reconciliation of multiple gene trees to create a robust species tree.

Species and gene trees are powerful tools to understand the evolutionary history of viruses and their hosts. While a species tree traces the evolutionary history of the species as a whole, gene trees trace the history of a specific gene, which may not necessarily mirror the species tree (Edwards, 2009; Swenson and El-Mabrouk, 2012). Species trees can be discordant with gene trees for several reasons, including gene duplication and loss, horizontal gene transfer, incomplete lineage sorting, and evolutionary rate variation among loci (Steenwyk et al., 2023; Maddison, 1997). Rate variation, in particular, can have a major effect on gene tree estimation due to substitutional saturation (multiple mutations occurring at the same site on a branch within the tree), which obscures nucleotide changes that have occurred, reducing the accuracy of phylogenetic trees inferred using such loci (Philippe et al., 2011; Philippe and Forterre, 1999; Stern and Andino, 2016). Collectively, these processes motivate species tree inference using data from numerous loci with varying rates of evolution, allowing discordance among individual gene trees with differential evolutionary rates to create a well-resolved species tree (Edwards, 2009). Using phylogenetic methods such as the multi-species coalescent, we can reconcile multiple discordant gene trees to construct a well-supported species tree for *Adenoviruses*, allowing us to accurately place *Anolis adenovirus 2* (Swenson and El-Mabrouk, 2012; Szllosi et al., 2015). Previous whole-genome sequences of *Adenovirus* have been used to infer phylogenetic trees that recover each genus as monophyletic with the following relationships: *Mastadenovirus* and *Barthadenovirus* resolved as sister taxa, followed by *Aviadenovirus*; *Siadenovirus* and *Testadenovirus* sister to each other, and *Ichtadenovirus* sister to the clade containing all other *Adenovirus* genera (Matsvay et al., 2022; Doszpoly et al., 2019; Miller et al., 2017). We predict that our phylogenetic tree will recover these relationships and that our two sampled strains of *Anolis adenovirus 2*, both drawn from the same host species, will be closely related to each other and to other squamate *Barthadenoviruses*.

Beyond reconstructing the evolutionary relationships among *Barthadenovirus* species, gene annotation can reveal how viral genomes have evolved with regard to gene content, genome organization, and rate of evolution per gene. We expect to find a suite of core genes present across all *Barthadenovirus* lineages, such as genes essential for viral replication or genes that code for structural components of the virus. We expect that these genes will be in the same relative order with minor rearrangements (Kulanayake and Tikoo, 2021; Matsvay et al., 2022; Davison et al., 2003). Additionally, we expect to find species-specific ORFs towards the ends of the genome: Previous research found that some species-specific genes coded for proteins that might be associated with host cell uptake (Matsvay et al., 2022; Farkas et al., 2008; Pénzes et al., 2020) Finally, analyzing the relative rate of evolution per gene may help us better understand how the virus is able to adapt to its host: In one bird-infecting *Barthadenovirus* DNA polymerase (DPOL) and the penton protein (CAPSP) were found to have experienced positive selection, which could be associated with adaptation to the host following a host-switching event (Matsvay et al., 2022). Integrating our gene annotation of *Anolis adenovirus 2* with comparative evolutionary analyses of other *Adenovirus* strains can allow us to distinguish conserved genomic features essential for viral function from rapidly-evolving regions that may help the virus adapt to its host.

In the present study, we had two primary objectives: First, we aimed to place *Anolis adenovirus 2* within the broader evolutionary context of *Adenoviruses*. Although members of *Barthadenovirus* are known to have undergone host switching, we hypothesized that *Anolis adenovirus 2* would be most closely related to other squamate *Barthadenoviruses*. Second, we used our genome annotation to investigate patterns of genome evolution and structure. We predicted that gene order would be conserved across *Barthadenovirus*, but individual genes would experience differential rates of evolution: Proteins that help the virus interact with and infect the host cell will likely experience faster rates of evolution than conserved structural and DNA replication proteins. To answer these questions, we assembled, annotated, and performed evolutionary genomic analyses of the whole-genome sequence of two *Anolis adenovirus 2* isolates found to be infecting two *Anolis distichus ravitergum* lizards. Our sequences facilitate the creation of a reconciled species tree from twenty-three orthologous genes, identify a set of core genes across *Adenovirus* and *Barthadenovirus* respectively, and calculate the relative rate of evolution of each gene. Taken together, this research provides new insight into the evolutionary history of *Anolis adenovirus 2*, the genome organization of *Adenovirus*, and the evolutionary forces shaping individual viral genes.

## 2 Methods

### 2.1 Laboratory Methods

The collection of *Barthadenovirus* samples selected for whole-genome sequencing was previously described by Ascher and colleagues (2013). Briefly, a small number of wild-caught *Anolis distichus ravitergum* lizards from the Dominican Republic became lethargic and sick shortly after importation and were humanely euthanized. Based on an analysis of a 300-base-pair region of the DNA polymerase gene amplified from DNA extracts of lizards’ gastrointestinal tract tissues, it was found that these anoles harbored a novel *Barthadenovirus* strain, Anolis adenovirus 2 (Ascher et al., 2013).

To generate whole-genome sequences of Anolis adenovirus 2, we used the extractions from Ascher et al. (2013) from two samples: Sample 8054 and sample 8158. For these samples, the previously extracted DNA was sheared to lengths of 500 base pairs using a Covaris S2 Focused Ultrasonicator (Covaris, Waltham, MA). Libraries were prepared using the NebNext Ultra II Library Prep Kit for Illumina (New England Biolabs, Ipswich, MA). Quantification was completed using the Qubit dsDNA High Sensitivity Kit (Thermo Fisher, Waltham, MA) and libraries were size-checked on a TapeStation instrument using D5000 tapes (Agilent, Santa Clara, CA). Once verified, paired-end (2 *×*150 bp) whole-genome sequencing was performed using an Illumina iSeq in GenerateFastq mode (2 *×* 150 base-pair length) (Illumina, San Diego, CA). We ran a total of three iSeq flow cells for each sample to generate enough viral DNA for a successful assembly.

### 2.2 Genome Assembly and Annotation

We used FastQC v0.11.9 to evaluate the quality of the generated raw sequence reads (Andrews, 2010). We removed adapter sequences and low-quality reads using Trimmomatic v0.39 (Bolger et al., 2014) and ran FastQC after trimming to verify that adapters were successfully removed and to assess the quality of the trimmed reads. Trimmed reads were used as input for SPAdes v3.15.5, which was run in metaviral mode (Prjibelski et al., 2020). From there, we conducted a BLAST search using BLAST v.2.14.0 to identify viral scaffolds. We used BWA 0.7.17-r1188 (Li and Durbin, 2009) to map the trimmed reads back onto the assembled viral scaffolds to calculate coverage and breadth statistics (Table 1). We used VGAS v.0.0.2020.07.22 (Zhang et al., 2019) to predict gene models and another BLAST search to functionally annotate the genes. Finally, we used the R package gggenes v.0.6.0 (Wilkins, 2023; R Core Team, 2026) to visualize the positions of these gene models along the virus.

**Table 1:** Sequencing effort: Summary statistics depicting assembled scaffold length, depth and breadth of coverage, and number of reads that were used to generate 8054 and 8158 viral assemblies.

| Anolis adenovirus 2 Sample ID | 8054 | 8158 |
| --- | --- | --- |
| Length | 37517 | 37535 |
| Depth of Coverage | 68.4067 | 62.942 |
| Breadth of Coverage | 100 | 100 |
| Number of Reads Mapping | 17344 | 15839 |
| Proportion of Reads Mapping to Reference | 0.19 | 0.15 |
| GC Content | 46.87% | 46.87% |

### 2.3 Phylogenetics

To infer our phylogeny, we downloaded a selection of related *Adenovirus* sequences from NCBI Genbank (Supplemental Table 1). We ran all sequences, including our two *Anolis adenovirus 2* sequences, through VGAS and BLAST v.0.0.2020.07.22 (Zhang et al., 2019) to functionally annotate them and used OrthoFinder v.2.5.5 (Emms and Kelly, 2019) to identify orthologous genes among sampled genomes. We selected the top 23 orthogroups, as these alignments contained at least 40% of all selected species. Furthermore, all of the top 23 orthogroups were associated with a functional annotation in our BLAST analysis. We excluded some orthogroups from OrthoFinder, as some of the previously-published Adenovirus sequences were only partial sequences (noted on tree) (Figure 3). The sequence alignments of the first 23 orthogroups were used as input for the program IQTree v.2.2.2.7 to make gene trees, which were then reconciled into a single species tree using ASTER-Pro3 (Zhang et al., 2025). White sturgeon adenovirus 1 was selected as the outgroup for the phylogeny due to its distant relationship to other *Adenovirus* sequences (Doszpoly et al., 2019). We used the R package ggtree v.4.0.5 for tree visualization (Yu et al., 2017; R Core Team, 2026).

To identify instances of potential host switching, we contrasted the reconciled virus tree topology described above with established phylogenies of host lineages. Specifically, we investigated four levels of potential incongruence. First, we checked whether the previously identified *Adenoviruses* genera were monophyletic. Second, as *Barthadenoviruses* are known to infect mammals, squamate reptiles, and birds, we checked for the monophyly of the virus strains infecting each of these vertebrate classes (Harrach, 2000). For instance, we asked if all mammal-infecting *Barthadenoviruses* share a common ancestor to the exclusion of all squamate and bird-infecting strains. Third, we compared the relationships among squamate-infecting *Barthadenoviruses* and the well-established relationships between these host species sampled in our tree. The consensus of many recent phylogenetic studies reflects the following host relationship: ((((Gila Monster, Snake), Bearded Dragon),Anole), Lacerta) (*reviewed in* Gable et al., 2023). Finally, we tested for potential host-switching involving either of our two sequenced isolates by asking if they represent sister lineages in our tree. Since both of our two sequenced isolates belong to the same adenovirus strain (*Anolis adenovirus 2*, we considered the possibility of host switching at this level highly unlikely.

### 2.4 Rate of Evolution

We predicted that genes in *Anolis adenovirus 2* would experience differential evolutionary pressures, and that genes that enable the virus to interact with the host cell would evolve faster. To measure whether certain genes were experiencing faster rates of evolution than others, we calculated the relative rate of evolution of each gene. We used the alignments of the previously-selected 23 orthogroups and concatenated them together into Nexus and Phylip alignment files using Concatenator v0.3.1, retaining a distinct partition for each gene (Vences et al., 2022). We then ran IQTree v.3.1.3 to calculate partition-specific rates which, due to our partitioning scheme, are equal to the relative rate of evolution for each gene (Wong et al., 2026).

### 2.5 Gene Arrangement

We predicted that the gene order of *Anolis adenovirus 2* would be highly conserved. To understand patterns of viral genome evolution and measure synteny across genes, we compared protein locations in *Anolis adenovirus 2* to selected *Barthadenovirus* sequences (*Barthadenovirus varani*, *Barthadenovirus zootocae*, *Bearded dragon adenovirus 1*, *Deer atadenovirus A*, *Lizard adenovirus 2*, *Tern adenovirus 1*, and *Psittacine adenovirus 3*). The squamate *Adenovirus* strains were selected based on the similarity of host species within the *Barthadenovirus* phylogeny, and the non-squamate species were selected as representatives of the mammal and avian *Barthadenovirus* genera (Figure 2). Protein sequences of 8054 and 8158 were reverse-complemented in R using dplyr for visualization purposes, and the visualization was created using gggenes v.0.6.0 (Wilkins, 2023; Wickham et al., 2019).

## 3 Results

### 3.1 Genome Assembly and Annotation

Sample 8054 and Sample 8158 assembled into scaffolds that were 37,517 and 37,535 base pairs long, respectively, which is within the expected genome size for *Barthadenovirus* sequences (Pénzes et al., 2014; Farkas et al., 2008; Athukorala et al., 2022) (Table 1). For both strains, the GC content was 46.87%. The depth and breadth of the genomes remain on par with previously-published *Barthadenovirus* genomes (Okoh et al., 2023; Matsvay et al., 2022). For both sequences, our annotation identified 36 potential genes; of these, 23 had a positive BLAST identity to a known viral protein (Figure 1). As with other *Barthadenovirus* sequences, essential genes (such as the DNA polymerase, fiber, spike protein, and capsid proteins) were located in the center of the genome with species-specific ORFs located towards the start and end of the genome (Lung et al., 2022; Pénzes et al., 2020) (Figure 4). Our two *Anolis adenovirus 2* genomes sequenced from samples 8054 and 8158 are extremely similar (>99%) to each other, likely due to the fact that they were sampled from two infected lizards collected from the same site at the same time (Ascher et al., 2013).

**Figure 1:**
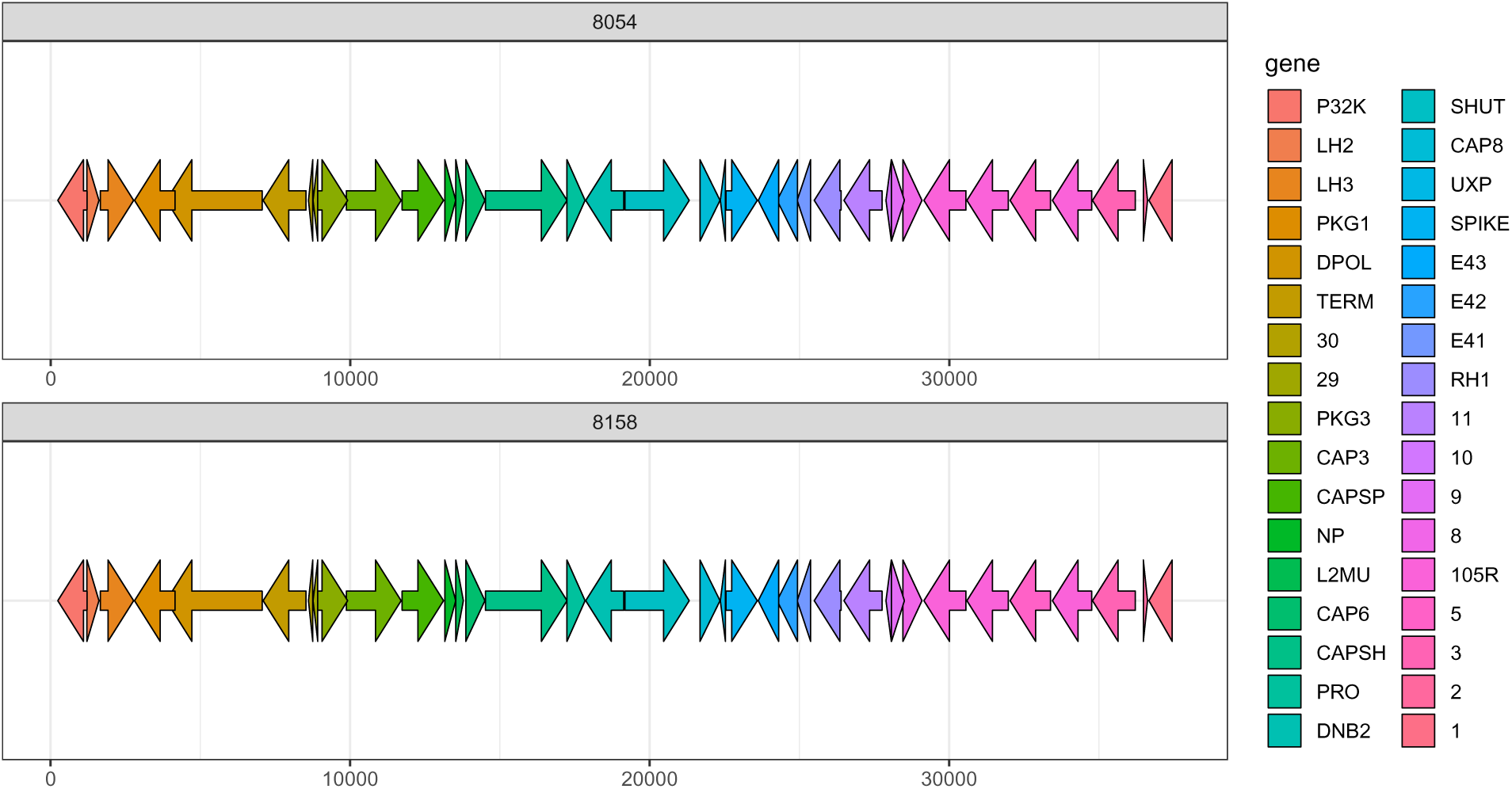
Gene Annotation for Samples 8054 and 8158: VGAS identified 36 potential ORFs from each sample. Of these 36, 22 had a positive BLAST identity to another known viral gene. A table linking short gene codes to full-length gene identities can be found in Table 2.

**Table 2:** Orthogroups and their partition-specific rates of evolution, with corresponding gene annotations.

| Orthogroup | Partition-Specific Rate | Gene Code | Gene ID |
| --- | --- | --- | --- |
| OG0000000 | 1.187 | SPIKE | Fiber protein |
| OG0000001 | 0.661 | CAP6 | Pre-protein VI |
| OG0000002 | 1.095 | CAPSP | Penton protein |
| OG0000003 | 1.898 | CAP3 | Pre-hexon-linking protein IIIa |
| OG0000004 | 0.734 | PKG3 | Packaging protein 3 |
| OG0000005 | 0.548 | TERM | Preterminal protein |
| OG0000006 | 0.794 | DPOL | DNA polymerase |
| OG0000007 | 0.751 | SHUT | Shutoff protein |
| OG0000008 | 0.750 | DNB2 | DNA-binding protein |
| OG0000009 | 0.783 | PRO | Protease |
| OG0000010 | 1.031 | CAPSH | Hexon protein |
| OG0000011 | 0.789 | CAP8 | Pre-hexon-linking protein VIII |
| OG0000012 | 0.471 | PKG1 | Packaging protein 1 |
| OG0000013 | 0.765 | E43 | Protein E4.3 |
| OG0000014 | 1.359 | NP | Pre-histone-like nucleoprotein |
| OG0000015 | 0.871 | RH1 | Protein RH1 |
| OG0000016 | 1.470 | E41 | Protein E4.1 |
| OG0000017 | 1.488 | E42 | Protein E4.2 |
| OG0000018 | 1.045 | LH3 | Hexon-interlacing protein LH3 |
| OG0000019 | 0.818 | L2MU | Late L2 mu core protein |
| OG0000020 | 1.384 | P32K | Structural protein p32K |
| OG0000021 | 1.454 | UXP | Probable U-exon protein |
| OG0000022 | 1.438 | LH2 | Protein LH2 |

### 3.2 Phylogenetic Analyses

We built a well-resolved phylogeny of all major *Adenovirus* clades and examined the phylogeny to test for the presence of host-switching at three levels: Across *Adenovirus*, within *Barthadenovirus*, and finally, within our two isolates of *Anolis adenovirus 2*. First, our *Adenovirus* species tree recapitulates all six genera (*Barthadenovirus*, *Mastadenovirus*, *Aviadenovirus*, *Siadenovirus*, *Testadenovirus*, and *Ichtadenovirus*) as monophyletic lineages. The species tree recovers *Barthadenovirus* and *Mastadenovirus* as sister clades, followed by *Aviadenovirus*, with *Siadenovirus* and *Testadenovirus* sister to each other. *Ichtadenovirus* is the outgroup and the most distantly related, as expected (Doszpoly et al., 2019) (Figure 2). Within *Barthadenovirus*, while our tree shows a likely reptilian origin of *Barthadenovirus*, the virus is shown to infect both birds and mammals. Additionally, our tree shows a deep set of relationships that are incongruent with the host phylogeny between *Bearded dragon adenovirus 1* and *Barthadenovirus zootocae* (Pincheira-Donoso et al., 2013). Finally, we expected that *Anolis adenovirus 2* would be nested within the squamate-infecting *Barthadenoviruses* based on its position in a tree inferred using a 300 base-pair region of the DNA polymerase gene (Ascher et al., 2013). Our species tree found that, as hypothesized, our two sequenced *Anolis adenovirus 2* samples (8054 and 8158) are closely related to other squamate *Barthadenoviruses* strains (Figure 2) and are sister taxa to each other (Pénzes et al., 2020, 2014; Farkas et al., 2008; Buck et al., 2024; Donald et al., 2021).

**Figure 2:**
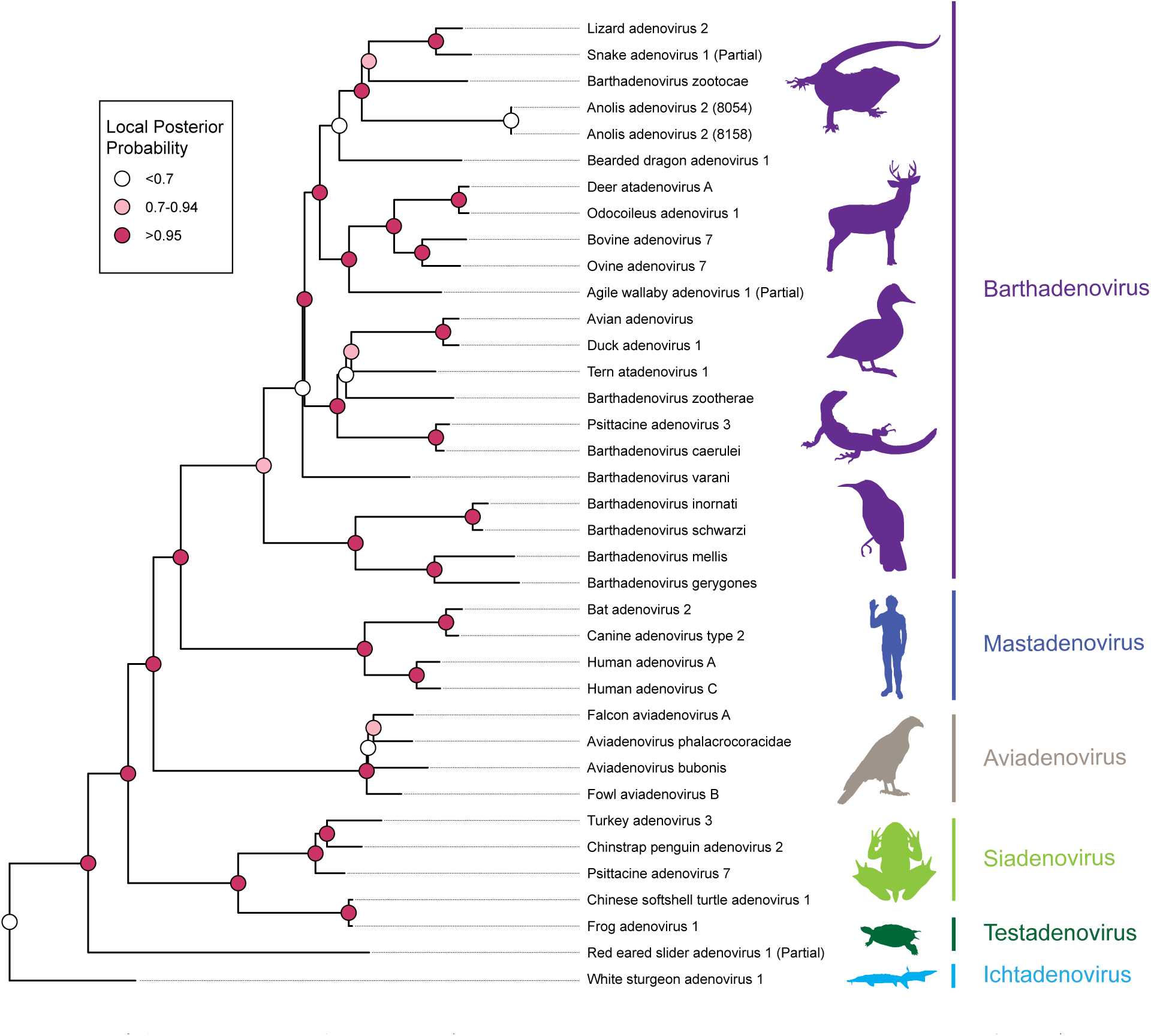
*Adenovirus* species tree: *Adenovirus* whole-genome species tree generated from Astral-Pro reconciliation of twenty-three orthogroup gene sequences with the colored circles at each node representing the local posterior probability values obtained from Astral-Pro. This species tree recovers all six previously-identified *Adenovirus* genera as monophyletic. Of specific interest within *Barthadenovirus* (purple), we find evidence of host-switching due to the discordant relationships of the virus between mammals, birds, and squamates within *Barthadenovirus*.

### 3.3 Orthologous Gene Analysis

We hypothesized that *Anolis adenovirus 2* would contain a “core” set of genes found in all *Adenovirus* sequences, as well as *Barthadenovirus*-and lineage-specific orthologues. We analyzed the presence or absence of the first 23 orthogroups, as these genes were all present in both 8054 and 8158 and had associated functional annotations (Table 2 and Figure 3). Our orthogroup analysis identified a “core” set of 13 genes that were present in all *Adenoviruses* across the entire phylogeny (excluding genomes with partial assemblies). Genes present in all whole-genome sequences of *Adenoviruses* analyzed included the penton protein (CAPSP), pre-penton protein VI (CAP6), Pre-hexon-linking protein IIIa (CAP3), Packaging protein 3 (PKG), preterminal protein (TERM), DNA polymerase (DPOL), shutoff protein (SHUT), DNA-binding protein (DNB2), protease (PRO), hexon protein (CAPSH), and the pre-hexon-linking protein VIII (CAP8) (Figure 3). In a majority of sequences, the fiber protein (SPIKE) and packaging protein 1 (PKG1) were present; we considered these to be included in the “core” genes. Previous analyses have classified anywhere between 9 and 16 core genes (Davison et al., 2003; Kulanayake and Tikoo, 2021). We observe that some proteins (LH2, LH3, P32K, E41, E42, and E43) appear to be present in a majority of the *Barthadenovirus* sequences (Figure 3), which align with previous results (Zheng et al., 2024; Gorman et al., 2005; Davison et al., 2003; Marabini et al., 2021; Both, 2002; Pantelic et al., 2008). Previous research has identified putative functions for some of these proteins: The LH3 protein is theorized to be associated with capsid assembly and the RH1 protein is predicted to help the virus infect host cells (Both, 2002; Pantelic et al., 2008). Finally, in *Anolis adenovirus 2*, we recover a homolog for the 105R gene, which is also present in Lizard-Adv-2 (host: *Heloderma horridum*), Snake-Adv-1 (host: *Pantherophis guttatus*), and *Barthadenovirus zootoca* (host: *Zootoca vivipara*), but was originally identified in tree shrew adenovirus 1 (*Mastadenovirus*; host: *Tupaia* spp.). (Pénzes et al., 2014; Farkas et al., 2008).

**Figure 3:**
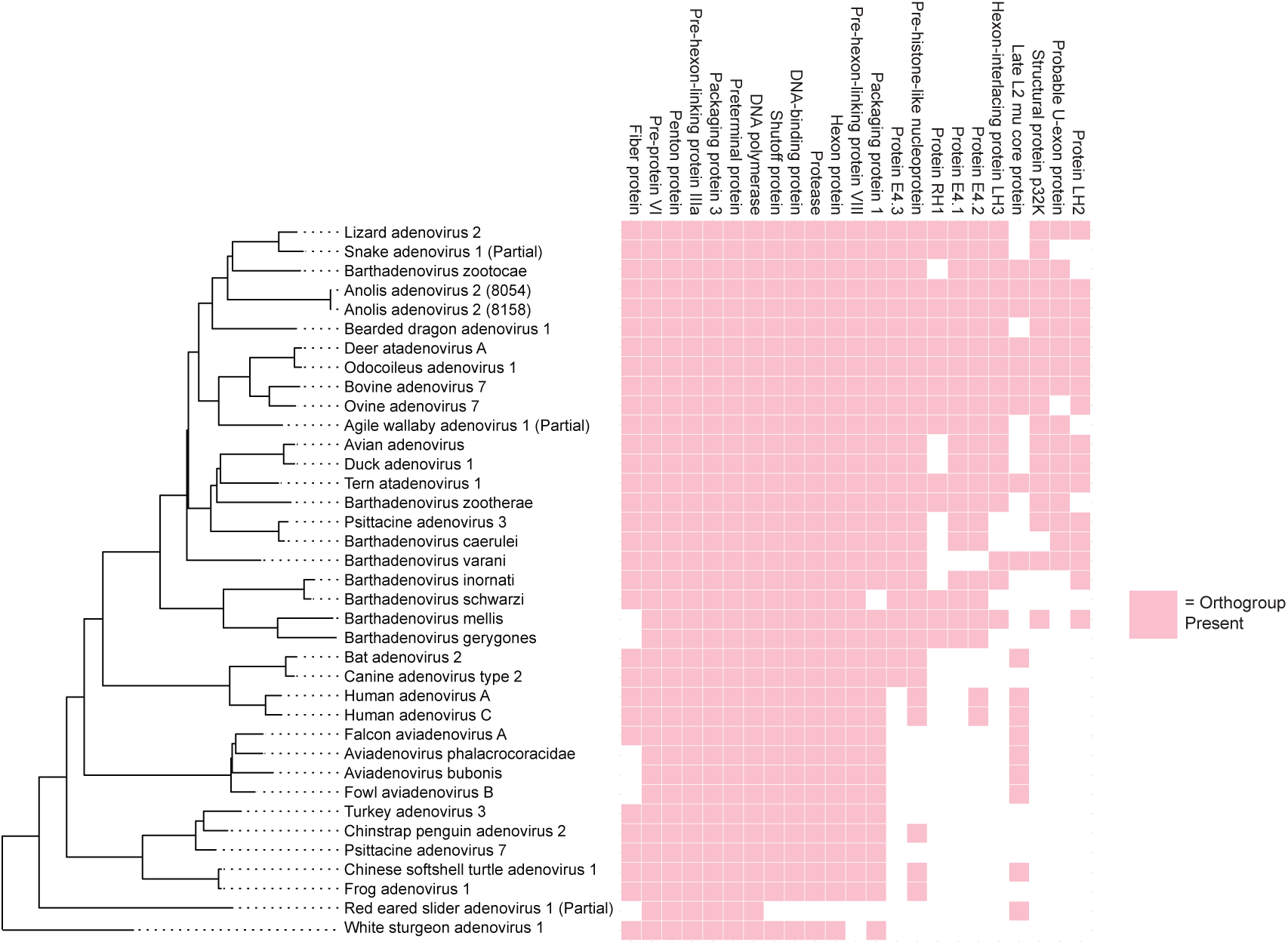
Orthogroup presence across *Adenovirus* strains: Our research identifies a “core” set of 13 genes that are present in a majority of the complete whole-genome sequences of *Adenovirus* (partial sequences are indicated). We count the fiber gene and the packaging protein within this core set of 13 genes. Additionally, we identify sdme genes that are present in a majority of the *Barthadenovirus* lineages, such as the E and L protein series.

### 3.4 Synteny

Gene order within all thirty-six identified ORFs in 8054 and 8158 shows the core group of proteins in the center of the viral genome, with species-specific ORFs towards the ends of the genome (Figure 1). This result is concordant with other previously-published *Adenovirus* phylogenies (Matsvay et al., 2022; Pénzes et al., 2020). Additionally, visualization of the conserved genes comparing *Anolis adenovirus 2* to a subset of *Barthadenovirus* revealed a genome that is largely stable with respect to gene order (Figure 4). Genes that are present across all *Barthadenovirus* have an identical conserved order except in *Deer atadenovirus A*, where the P32K protein and LH3 protein have appeared to have switched positions (Figure 4). There are slight variations in numbers and locations of genes, particularly on both ends of the genome, including the LH2, LH3 E4.1, E4.2, and E4.3 genes, across *Barthadenovirus*. Additionally, in *Anolis adenovirus 2*, we recover a homologue for the 105R gene, which has been found in a subset of *Barthadenovirus* strains; its location in *Anolis adenovirus 2* is similarly located on the end of the genome, near the species-specific ORFs (Figure 1) (Kulanayake and Tikoo, 2021; Pénzes et al., 2020). Altogether, the major genes involved in DNA replication and viral assembly present in *Anolis adenovirus 2* and other reptilian adenoviruses (*Barthadenovirus varani*, *Barthadenovirus zootocae*, *Bearded dragon adenovirus 1*, and *Lizard adenoivrus 2*) are all in an identical, conserved order.

**Figure 4:**
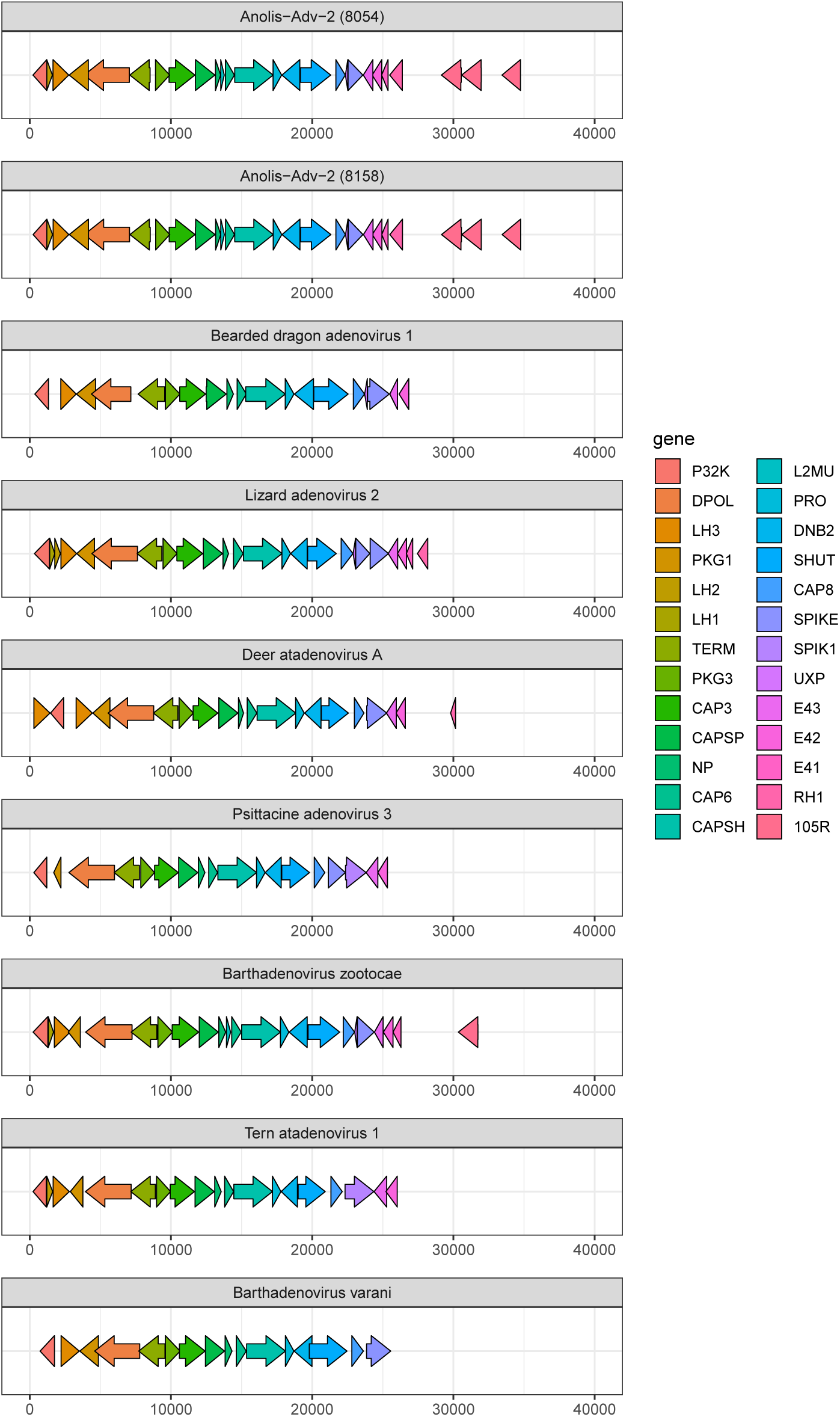
Gene order across *Adenovirus* strains: We found that gene order is largely conserved across *Barthadenovirus* and that only minor rearrangements have occurred throughout evolutionary history.

### 3.5 Rate of Evolution

We expected that genes across *Barthadenovirus* would experience different rates of evolution due to varying levels of purifying selection. We measured the rates of evolution by calculating the partition-specific rate for each gene in IQTree, where values greater than 1 indicate that the gene is evolving slower than average and values less than 1 indicate that the gene is evolving faster than average. We found that the fastest-evolving gene was the pre-hexon linking protein IIIa (CAP3), with a partition-specific rate of 1.898, nearly twice the genome-wide average. Other quickly-evolving genes include E4.2 (1.488), E4.1 (1.47), and the U-exon protein, UXP (1.454). In contrast, the slowest-evolving genes include the Packaging Protein 1 (PKG1) (0.471), the preterminal protein (TERM) (0.548), the pre-protein VI (CAP6) (0.548). Structural genes like the hexon (CAPSH) (1.031), penton (CAPSP) (1.095), and fiber (SPIKE) (1.187) proteins have rates of evolution that are closer to the genome average (Table 2). Finally, the DNA polymerase gene (DPOL), commonly used for constructing *Adenovirus* phylogenies, is under a slower rate of evolution than the genome average (0.794) (Wellehan et al., 2004b).

## 4 Discussion

It has been previously established that host-switching is relatively common among *Adenovirus* genera, particularly in *Barthadenovirus*, with widespread discordance between the host and virus phylogenies (Athukorala et al., 2022; Harrach, 2000). At the generic level, our phylogeny recovers all six *Adenovirus* genera as monophyletic, and this result, including their specific branching order, is concordant with previous studies (Mukai et al., 2019; Davison et al., 2003; Harrach et al., 2019; Salzmann et al., 2021; Matsvay et al., 2022; Zheng et al., 2023; Tian et al., 2025; Athukorala et al., 2022; Doszpoly et al., 2019). In *Barthadenovirus*, we observe several potential instances of host switching: Our tree supports the previously-hypothesized sqamate reptile origin of *Barthadenovirus* (Wellehan et al., 2004b) and find *Barthadenovirus* strains infecting both birds and mammals, particularly ruminants (Harrach, 2000; Kaján et al., 2020; Athukorala et al., 2022) (Figure 2). This distribution of hosts is likely the result of past host-switching from squamate reptiles into these novel host lineages. We additionally observe a second potential ancestral host-switching event within squamate *Barthadenoviruses*: our viral phylogeny is incongruent with their host species tree (Gable et al., 2023). Specifically, the phylogenetic positions of *Barthadenovirus zootocae* and *Bearded dragon adenovirus 1* are discordant with recent trees inferred for these groups (Gable et al., 2023). Finally, our phylogenetic tree recovers our two novel whole-genome sequences of *Anolis adenovirus 2* as sister taxa. While they are solidly placed within the squamate *Barthadenoviruses*, there is no signature of host-switching between these two taxa and other lineages, which aligns with previous findings (Ascher et al., 2013) (Figure 2).

Previous analyses of adenoviruses infecting *Anolis* have suggested that host switching may be occurring among *Anolis* host species (Prado-Irwin et al., 2018). Despite this, previous research into *Anolis adenovirus 2* based on a 300 base-pair region of the DNA polymerase gene placed this strain within *Barthadenovirus*, closely related to other squamate-infecting strains, rather than other clades of *Adenovirus* (Ascher et al., 2013). Our phylogeny does not suggest that *Anolis adenovirus 2* has experienced a large-scale host switching event, as these two isolates are identified from the same strain and same subspecies, albeit in two distinct

*Anolis distichus ravitergum* individuals (Figure 2). However, our species tree lacks the fine-scale sampling of viruses infecting *Anolis* lizard that would be needed to resolve whether host-switching is definitively occurring. We have made substantial efforts, but have been to-date unsuccessful, in sequencing the genomes of other anole-infecting *Adenoviruses* (data not shown). Whole-genome sequencing of other previously-published *Barthadenovirus* strains found to infect *Anolis*, as well as sampling of novel, wild taxa across *Anolis* will be necessary to fully understand whether host-switching is happening at the species level (Prado-Irwin et al., 2018; Ascher et al., 2013). Given the fact that there are many sampling gaps in *Barthadenovirus*, let alone the more than four hundred species of *Anolis*, screening an expanded diversity of potential host taxa for this virus would help us to identify understand any patterns underlying the host shifts that have occurred as *Barthadenovirus* evolves.

Our analyses identified a set of 13 proteins present in all complete whole-genome sequences of *Adenovirus*, which largely included the proteins necessary for capsid formation and viral replication (Sohaimi and Hair-Bejo, 2021; Kulanayake and Tikoo, 2021) (Figure 3, Table 2). We also identified the fiber protein (SPIKE) in a majority of the represented sequences (Figure 3). The fiber protein is known to have high sequence divergence among adenoviruses, particularly fowl aviadenoviruses (Sohaimi and Hair-Bejo, 2021). We did not detect an ortholog of fiber protein among bird-infecting *Barthadenovirus* strains, however this could be due to this high divergence. We identify the packaging protein in all strains except for *Barthadenovirus schwarzi*. The eleven unidentified ORFs towards both far ends of *Anolis adenovirus 2* are not uncommon in *Adenovirus* strains (Matsvay et al., 2022; Zheng et al., 2023; Athukorala et al., 2022; Kohl et al., 2012) but further research into *Barthadenovirus*, including functional annotation studies, are necessary to understand what, if any, proteins are produced from these ORFs in Anolis adenovirus 2. The presence of the gene 105R in our viruses adds another reptilian taxon in which this gene has been identified (Farkas et al., 2008; Pénzes et al., 2014). Gene 105R was originally described in mammals and has otherwise been reported so far only in squamate-infecting *Barthadenovirus*. Our study identifies 105R in *Barthadenovirus zootocae*, which also infects squamate (*Zootoca vivipara*) (Figure 3). Its patchy distribution, together with the sister relationship between *Mastadenovirus* and *Barthadenovirus*, suggests that the evolutionary history of 105R warrants further investigation and may provide a useful system for exploring hypotheses of host switching and viral speciation between the two clades (Schöndorf et al., 2003). Our results suggest that genome structure is largely conserved across *Adenovirus*, with genes across *Anolis adenovirus 2* appearing in the same order as other strains of squamate *Barthadenovirus* (Figure 4) (Pénzes et al., 2014, 2020).

Our synteny results mirror the results from previously-published studies with regard to the order of identified genes (Matsvay et al., 2022; Pénzes et al., 2020, 2014). Altogether, there is a high conservation of gene order across *Adenovirus* that aligns with previous studies, which found that even adenoviruses from distantly related species such as between birds and humans had extreme similarity with regard to synteny (Pitcovski et al., 1998; Vrati et al., 1996).

As predicted, we find differential rates of evolution per gene in *Adenovirus* (Table 2). Overall trends from our evolutionary rate analysis show that genes that are involved with viral replication, such as DNA polymerase (DPOL), the shutoff protein (SHUT), DNA binding protein (DNB2), preterminal protein (TERM) -whose activity is closely dependent on DNA polymerase, packaging protein (PKG1), and protease (PRO), experience slower rates of evolution than average (Table 2) (Davison et al., 2003; Webster et al., 1997; Greber et al., 1996; Condezo et al., 2015). In contrast, proteins coding for the major structural components of the virus, such as the fiber protein (SPIKE), penton protein (CAPSP), and hexon protein (CAPSH) have values slightly above 1 (Table 2). These proteins mediate host-cell attachment, receptor binding, cell entry, and are major targets of the host immune response, which makes them subject to greater evolutionary diversification than the highly-conserved replication proteins (Davison et al., 2003; Robinson et al., 2011; Kulanayake and Tikoo, 2021). Two of three E4 series proteins (E4.1 and E4.2), which have been shown to modulate viral and cellular growth and gene expression, and are shown to be related to the virus’s ability to interact with the host, are under quick rates of evolution (Täuber and Dobner, 2001; Evans and Hearing, 2003). Interestingly, the fastest-evolving protein, pre-hexon linking protein IIIa (CAP3), is a structural protein that is known to help link the viral capsid together and help with viral packaging (Ma and Hearing, 2011; Weber, 1995). As such, the pre-hexon linking protein IIIa (CAP3) represents a notable exception to the general pattern observed in other structural proteins, suggesting that not all capsid-associated proteins are constrained. Taken together, functional role is a major determinant of evolutionary rate in adenoviruses. Proteins involved in replication are usually highly conserved, likely the result of purifying selection maintaining the core viral function of these genes. In contrast, genes involved in virion structure and host interactions exhibit faster rates of evolution due to either reduced functional constraint, or more likely, as the result of repeated bouts of antagonistic co-evolution between these viral proteins and host defenses.

Many previous *Adenovirus* phylogenies, including our own, have been based on the DNA polymerase gene (DPOL) due to its conservation across all *Adenovirus* strains and ease of amplification using widely available primer sets (Wellehan et al., 2004b; Prado-Irwin et al., 2018; Ascher et al., 2013; Henriques et al., 2014; Kohl et al., 2012). Furthermore, this gene appears to occupy an evolutionary sweet spot as it evolves fast enough to detect sequence differences between relatively closely related strains, yet slow enough to largely avoid mutational saturation when comparing more distantly related strains, such as in the case of the *Barthadenovirus* squamate to ruminant host-switching event (Yang and Rannala, 2012; Kapli et al., 2020). Our analysis of relative evolutionary rates supports DNA polymerase as an excellent choice for *Adenovirus* phylogenies due to its presence across all *Adenovirus* sequences and its slower-than-average rate of evolution. Our analysis also helps identify additional slowly-evolving genes that could be targeted for future primer design to build robust phylogenies, particularly for deeper relationships. Both preterminal protein (TERM) and pre-protein VI (CAP6) are present in all sampled *Adenoviruses* and have a slow rate of evolution; the slowest-evolving protein, packaging protein 1 (PKG1), is present in all sequences studied with the exception of *Barthadenovirus schwarzi* (Table 2; Figure 3). The identification of these conserved, slowly evolving genes supports future studies aimed at resolving deep evolutionary relationships, host-switching events, and the taxonomy of *Adenovirus*.

Our high-quality genome assembly of *Anolis adenovirus 2* is the first whole-genome sequence of an *Adenovirus* from an *Anolis* lizard. These data enabled our investigation of the broader phylogenetic patterns in *Barthadenovirus* and supported previous observations of host switching at multiple evolutionary scales, highlighting the virus’s propensity and ability to switch between both distantly and closely related hosts. Our annotation analyses show that, despite a significant amount of conservation with respect to gene order, genes that are associated with interactions with the host cell experience faster rates of evolution, potentially facilitating adaptation to novel host species. Altogether, our assemblies add to an expanding number of *Adenovirus* whole-genome sequences, enabling future investigation of the relative roles of viral host-switching and coevolution across *Adenovirus*.

## Declaration of generative AI and AI-assisted technologies in the manuscript preparation process

During the preparation of this work, the authors used ChatGPT for help with coding, particularly with the creation of the phylogenetic tree and heat map in R, and minor language revisions during the writing process. The authors evaluated the output of ChatGPT’s code and grammar suggestions, reviewed and edited the content, and take full responsibility for the content of the published article.

## Data Availability Statement

Data are available at NCBI under BioProject PRJNA1476751, BioSamples SAMN60730727 and SAMN60730726. These data are currently embargoed but will be released upon publication. Processed analyses files are similarly embargoed to the public and will be released upon publication. These files are available to reviewers at this private link: https://dataverse.harvard.edu/previewurl.xhtml?token=c64e79f0-80e9-4d13-8f6a-7fcbd2f3f6bc. Scripts created to perform the analysis can be found at: https://github.com/chfal/virus_analyses.

## Acknowledgments and Funding Sources

This work was supported by NSF grants DGE-2152059 and DBI-2217717 to AJG, and GRFP-2240918 to CHF. We thank Sofia Prado-Irwin, Regan Kenia, Sashoya Dougan, Andrew Ebenezer, Alexis Winters, MacKenna Durbin, and Adesupo Adetowubo for their assistance in the laboratory. We thank the Coriell Institute for Medical Research for assistance with sonication and we thank Rutgers OARC for access to computational resources.

---

**Supplemental Table 1:**
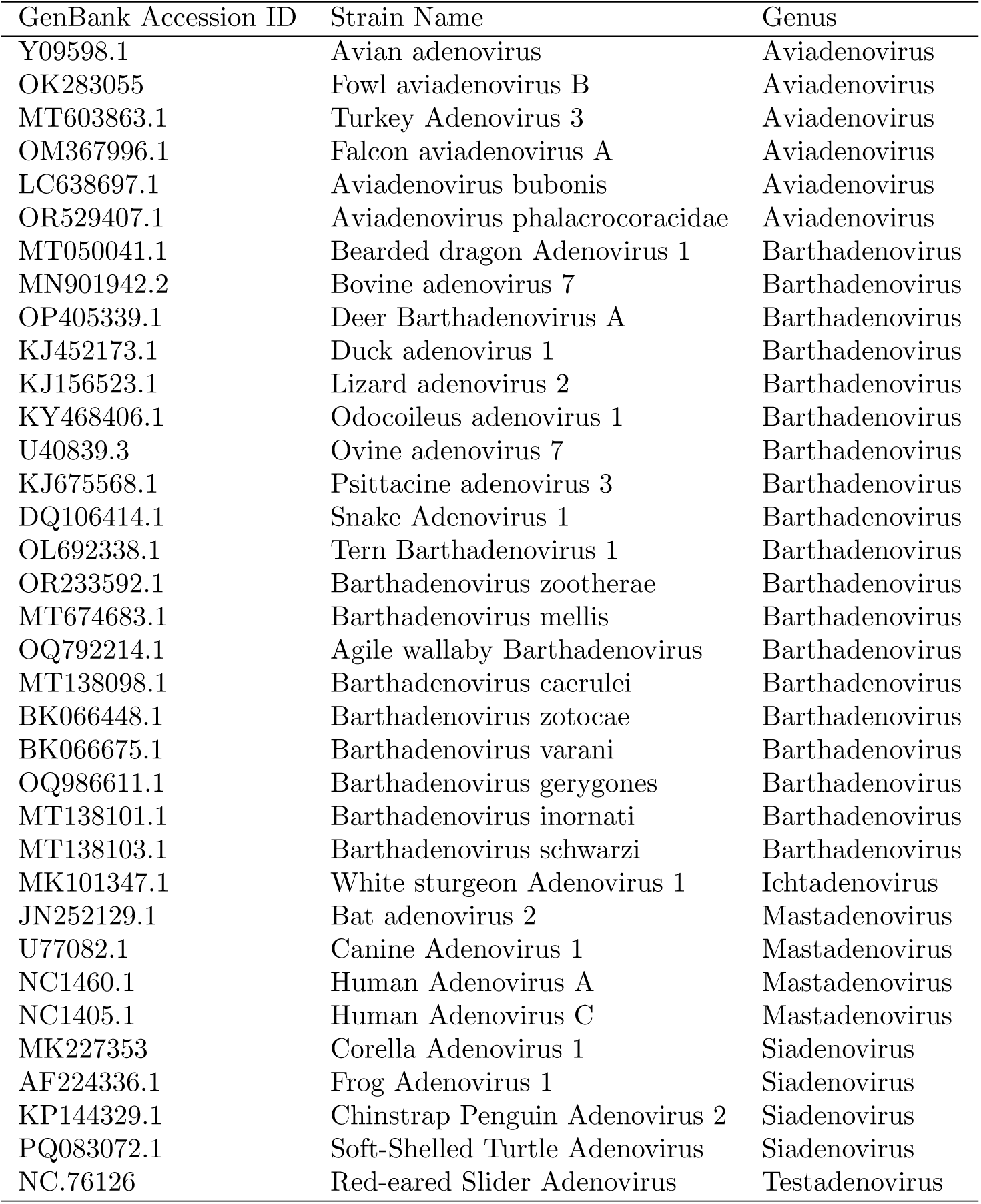
Accession numbers of *Adenovirus* strains downloaded from GenBank for our phylogenetic analyses and their corresponding names and genera.

